# Comprehensive and empirical evaluation of machine learning algorithms for LC retention time prediction

**DOI:** 10.1101/259168

**Authors:** Robbin Bouwmeester, Lennart Martens, Sven Degroeve

## Abstract

Liquid chromatography is a core component of almost all mass spectrometric analyses of (bio)molecules. Because of the high-throughput nature of mass spectrometric analyses, the interpretation of these chromatographic data increasingly relies on informatics solutions that attempt to predict an analyte’s retention time. The key components of such predictive algorithms are the features these are supplies with, and the actual machine learning algorithm used to fit the model parameters.

We here therefore evaluate the performance of seven machine learning algorithms on 36 distinct metabolomics data sets, using two distinct feature sets. Interestingly, the results show that no single learning algorithm performs optimally for all data sets, with different algorithm types achieving top performance for different types of analytes or different protocols. Our results can thus be used to find an optimal retention time prediction algorithm for specific analytes or protocols. Importantly, however, our results also show that blending different types of models together decreases the error on outliers, indicating that the combination of several approaches holds substantial promise for the development of more generic, high-performing algorithms.

## 1 Introduction

Mass Spectrometry (MS) coupled to Liquid Chromatography (LC) is a popular technique for the high-throughput analysis of the metabolome and lipidome, as it separates analytes based on their physicochemical properties [22, 4]. This is important because analytes compete for charges during ionization, leading to a strong bias against low abundant analytes. [34]. Moreover, LC is also capable of separating isobaric analytes, which plays a particularly important role in lipidomics [1].

LC-coupling thus provides analyte information that is complementary to the mass-over-charge (m/z) measurement of the MS. In many cases, high performance LC is used, where a solvent (the mobile phase) is pumped over a column (the stationary phase) under high pressure. The time an analyte takes to travel across the column is then determined by the degree of analyte interaction with the stationary and mobile phases, respectively, and is called the retention time (*t*_*R*_).

However, this LC retention time is usually not incorporated in the down-stream analysis, because it is either unknown *a priori*, or only known *a priori* for a very specific experimental setup. To fill this knowledge gap, researches typically use regression models to predict retention times for known metabolite structures. These *t*_*R*_ spredictions are based on a mapping between known structural, chemical and physical descriptors (or features) of the metabolites with their experimentally observed retention times (the targets). These predictions have previously been applied in both targeted and untargeted MS experiments to aid the analysis of lipids [1], metabolites [34, 11, 33] and peptides [19, 28, 16, 25].

For targeted MS the predicted *t*_*R*_ has been applied to reduce the number of experiments needed to study specific analytes of interest [3], while in untargeted MS these predictions have been used to differentiate between isobaric lipids [1], to filter false identifications for small metabolites (*<* 400 Da) [11], and to increase the number of confidently identified peptides [30].

As shown in the literature, retention time prediction can be a useful source of information for an MS experiment, but due to the many different experimental setups used across labs, the modeling procedure is not trivial [29, 5]. This because differences in setup significantly influence retention times, thus resulting in non-transferable knowledge between setups. To alleviate this lack of transferability of models, calibration approaches between different setups have been published, but these are only applicable to a limited set of metabolites [29] or setups [5]. As a result, researchers generally fit a new model for every setup, which requires substantial effort and data each time. Moreover, multiple modelling decisions have to be made for each new model that is trained, which leads to substantial heterogeneity and possible suboptimal modelling choices.

The regression model used for *t*_*R*_ prediction is generally fitted using a machine learning algorithm, and there is a large variety of algorithms to choose from [12]. For *t*_*R*_ prediction, the support vector regression model is the most popular option [27, 25, 19, 14, 18, 8, 20, 1], but other types of machine learning algorithms have been used as well, including linear regression, neural networks and random forest [34, 21, 7]. The choice of the algorithm is often guided by existing experience of the researcher with particular machine learning algorithms, which means that suboptimal approaches are often chosen by default. Indeed, many *t*_*R*_ prediction publications do not even justify the choice for their algorithm.

The field would therefore benefit from a comprehensive overview of the performance of different machine learning algorithms for *t*_*R*_ prediction, tested on a variety of experimental setups. We here therefore evaluate the performance of seven machine learning algorithms, applied to 36 distinct metabolomics data sets, using both a comprehensive feature set, as well as a minimal feature set. Our results show that the choice of one machine learning algorithm over another can significantly influence the performance, and that this choice is dependent on the characteristics of the data set.

## 2 Experimental section

### 2.1 Machine learning algorithms

A diverse set of seven machine learning algorithms is evaluated for their ability to compute accurate *t*_*R*_ prediction models from relatively limited amounts of source data. These algorithms all produce a regression model that relies on molecular descriptors (features) to predict the *t*_*R*_ (target). These algorithms were selected based on the fundamental differences in their prediction models, and on their popularity within both the LC and machine learning community.

Of these seven algorithms, two employ linear models, four employ non-linear models, and one features a hyperparameter that allows it to employ either a linear or a non-linear model. Linear models assume the relation between features and target to be linear and as such cannot capture more complex relations. However these models are typically much more robust towards overfitting of the data. Two popular linear models were chosen, which differ mainly in the way the linear model parameters are fitted to the data. These are Bayesian Ridge Regression (BRR) [15], and Least Absolute Shrinkage and Selection Operator regression (LASSO) [31].

In contrast, non-linear models are able to model more complex relationships between features and target, but this flexibility comes at the cost of an increased risk of overfitting, especially on small data sets with many noisy features. Four models were chosen here: a feedforward Artificial Neural Network (ANN), and three decision tree ensemble models that differ in the way the decision trees are constructed: through Adaptive Boosting (AB) [13], Gradient Boosting (GB) [23] or bagging in a Random Forest (RF) [6].

The final selected model is a Support Vector Regression (SVR) [10], which can take either the form of a linear SVR (LSVR) if no kernel is applied, or the form of a non-linear SVR (SVR) if a Radial Basis Function (RBF) kernel is applied.

### 2.2 Datasets

The machine learning models were fitted on 36 publicly available LC-MS datasets (Table S-1): 19 were obtained from MoNA (http://mona.fiehnlab.ucdavis.edu/), 16 from PredRet [29], and one from **(author?)** [1]. Each data set contained measurements for at least 40 unique analytes. These 36 datasets were acquired in different labs, using different experimental setups. Across all datasets, 8305 molecules were observed, of which 6759 are unique. These molecules cover a broad range of masses and chemical compounds: from 59.07 Da to 2406.65 Da (see Figure S-1 for the mass distribution), and from acetamide to lipids. Duplicated molecules in the same dataset were removed based on their SMILES representation [32].

### 2.3 Molecular descriptors (features)

RDKIT [17] is used to convert the SMILES representation of the molecules to 196 features (see Table S-2 for a complete list). A selection of 151 features is made form this list, and these 151 are used for training. Selection of these 151 features is based on a filter for standard deviation of a feature across the different molecules (*stdev* ¿ 0.01), and on Pearson correlation between features (*r*^2^ ¡ 0.96).

In addition, a minimal subset of eleven features (obtained from **(author?)** [1]) was selected to evaluate the potential for overfitting of the larger feature set. These eleven features include an estimate of the hydrophobicity using the octanol/water partition coefficient (MolLogP, slogp VSA1 and slogp VSA2), molar refractivity (SMR), estimated surface area (LabuteASA and TPSA), average molecular weight (AMW), polarizability based on molar refractivity (smr VSA1 and smr VSA2) and electrostatic interactions (peoe VSA1 and peoe VSA2).

### 2.4 Performance metrics

The generalization performance of a fitted regression model is evaluated using three different performance metrics: Mean Absolute Error (MAE), Median Absolute Error (MedAE), and Pearson correlation (*r*).

Let *ŷ* be the predictions of the model and *y* the experimentally observed retention times for all molecules *n* in a dataset; then the MAE is calculated using the following equation:

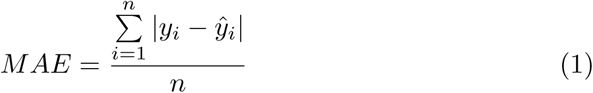

While MAE can give a good indication of performance, it is sensitive to outliers. For this reason the related but more robust Median Absolute Error is here used as the main metric:

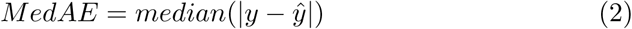

The MAE or MedianAE can be hard to compare between different classes of problems, especially when the error depends on the range of the elution times. Therefore, in addition to the previous two metrics, the Pearson correlation is calculated as well:

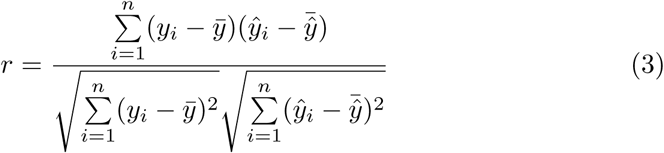

Even though the MAE, MedianAE, and correlation can be calculated for each datasets, a direct comparison between these metrics across different datasets is not possible. For instance, a prediction error of ten seconds will have more impact when the run time is 129 seconds (as is the case for dataset RIKEN) as compared to a run time of 4089 seconds (as is the case for dataset Taguchi). This can be alleviated by normalization; specifically by dividing by the error by the retention time of the last detected analyte:

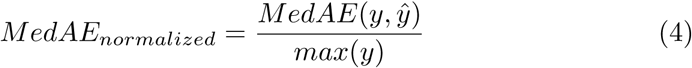

### 2.5 Learning Curves

As *t*_*R*_ prediction is specific to an experimental condition, the number of annotated molecules (the training set) available for fitting (or training) a regression model is typically limited. It therefore is important to investigate how different regression models perform for different training set sizes.

In order to compute the learning curves (which plot performance *versus* data set size), sufficiently large datasets are necessary. We therefore used datasets that contained at least 320 unique molecules for this specific analysis. The datasets that meet this requirement are Eawag XBridgeC18, FEM long, RIKEN, Stravs and Taguchi 12. To investigate the reproducibility of the obtained results, each experiment is repeated ten times with different random seeds.

For each dataset, 160 unique molecules are selected at random to form the training set. The remaining molecules in that dataset constitute the test set, which is used to evaluate model performance. The training set of 160 molecules is then sampled for molecule sets of increasing sizes. The smallest such training subset contains 20 molecules, sampled at random without replacement. Regression models are optimized and fitted on this training set and the resulting models are evaluated on the test set. In the next iteration, another 20 molecules are sampled at random without replacement from the remaining 140 selected molecules, these are added to the training set. A model is again optimized on this larger training set, and evaluated on the same original test set. This procedure is repeated until the training set contains 160 molecules.

### 2.6 Algorithm Performance Evaluation

The generalization performance of each algorithms is evaluated using 10-fold Cross-Validation (10CV), in which all unique molecules are randomly assigned to one of ten subsets of equal size, and each subset is in turn assigned as the test set, while the remaining nine subsets are used as the training set. This process is repeated ten times, such that each subset is used once for testing.

### 2.7 Hyperparameter optimization

Learning algorithms have hyperparameters that are used to fit the model parameters, and these hyperparameters need to be set by the user before fitting the data. Here, the hyperparameters of the learning algorithms are optimized using values that are randomly drawn from a prespecified distribution (see Code Listing S-1 for the definition of these distributions). Random optimization is generally able to find the optimal parameters significantly faster than a grid search, because it is hypothesized that it samples more of the parameter space for the same amount of iterations [2]. For each machine learning algorithm, a total of 100 randomly selected hyperparameter sets are evaluated using a 10CV. The hyperparameter set with the best mean absolute error using Cross-Validation (CV) is then used for fitting the complete training set.

### 2.8 Code and dataset

The code to generate the models implements the following libraries: scikit-learn V0.18.0 [26], Pandas V0.19.021 [24], RDKIT V2016.03.1 [17] and XGBoost V0.4 [9]. The code used to generate the regression models, to make predictions, and to produce the figures is available at: https://github.com/RobbinBouwmeester/tRPredictionOverview

## 3 Results and discussion

SVR models are one of the most frequently applied algorithms for LC *t*_*R*_ prediction [27, 25, 19, 14, 18, 8, 20, 1]. Even though a large variety of machine learning algorithms is available to researchers to train a *t*_*R*_ prediction model, there can be significant performance differences between these. In this section we therefore evaluate the performance of different machine learning algorithms on different *t*_*R*_ prediction tasks.

### 3.1 Prediction performance versus training set size

First, the ability of different learning algorithms to generalize training sets with different sizes is investigated. Figure 1 shows learning curves for each of the regression models for the five datasets with more than 320 molecules. As expected, the learning curves show that all regression models benefit from larger training sets. For the five datasets used here, the largest generalization performance gain is typically observed when doubling the dataset size from 20 to 40 training molecules. Still, most models show optimal performance when trained on the largest sample size of 160 molecules, with adaptive boosting as the notable exception. This because adaptive boosting tends to overfit on the larger training sets, which is particularly evident for the RIKEN dataset.

**Figure 1:**
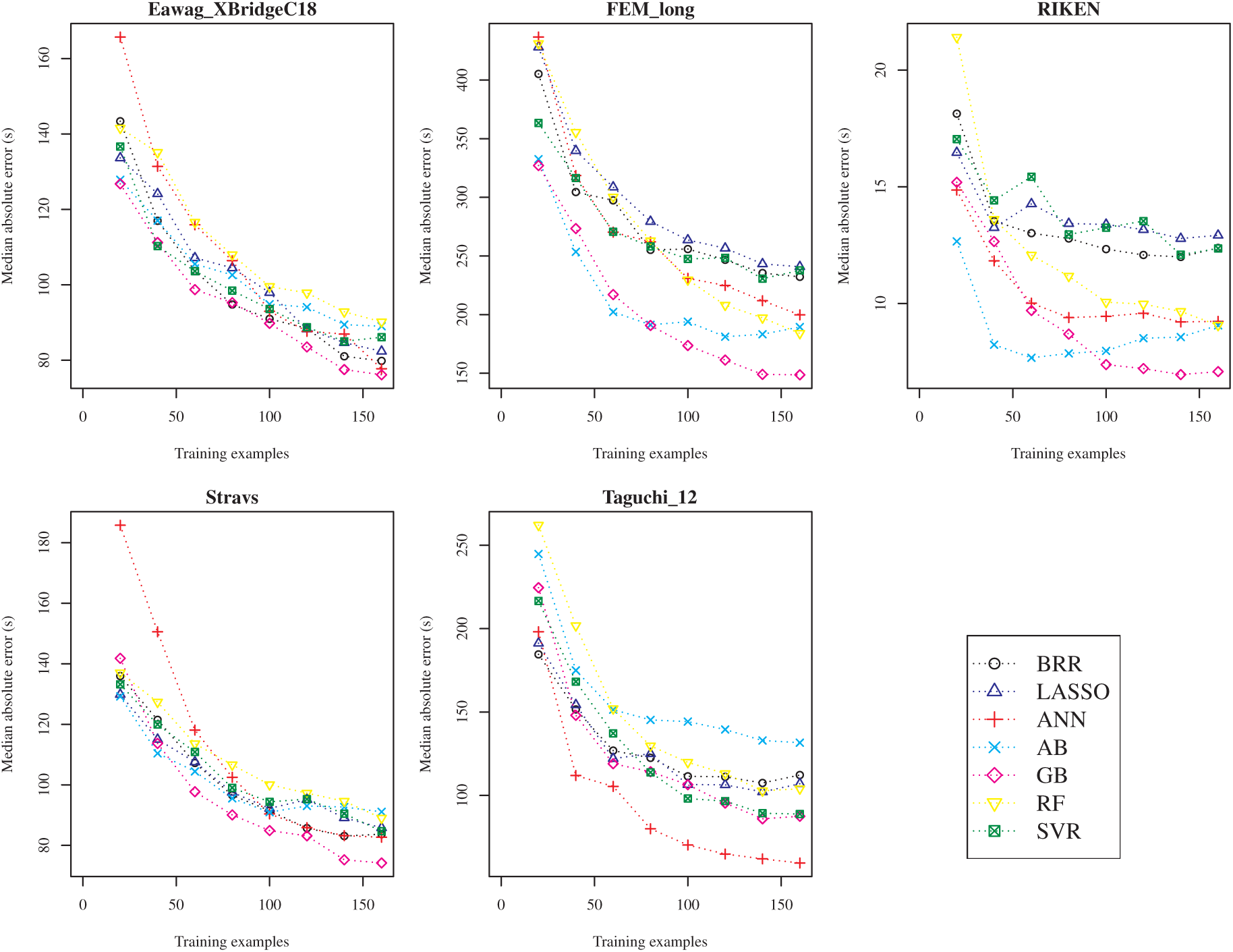
The learning curves for five datasets that have at least 320 training examples. The median absolute error (y axis) is plotted against a specific number of training examples (x axis). Training and testing for the points in the learning curve is repeated ten times, and the mean is plotted.

Although no clear performance difference between the linear and non-linear prediction models can be observed for the Eawag XBridgeC18 and Stravs datasets, there is a large performance increase for non-linear models for the FEM long, RIKEN and Taguchi datasets as training sets increase in size. For these latter datasets the GB algorithm clearly performs best overall, except for the Taguchi dataset where ANN clearly outperforms the other algorithms. This shows that the optimal algorithm is dataset dependent, and that boosting a decision tree forest (GB) performs significantly better than bagging a forest (RF) for *t*_*R*_ prediction. The same results are seen when plotting the mean absolute error, and the Pearson correlation (see Figure S-2 and S-3, respectively).

### 3.2 Prediction performance of the different algorithms on all datasets

In this section performance differences between the different regression models are evaluated on all 36 datasets. For each dataset, all unique molecules in that dataset are used in a 10CV approach to assess the generalization performance. The detailed CV results for mean absolute error, median absolute error, and Pearson correlation can be found in Tables S-3, S-4, and S-5, respectively.

Figure 2(a) shows the number of times an algorithm had the lowest median absolute error on any of the 36 datasets. GB again clearly stands out here, as it is the best performing algorithm for thirteen datasets. Linear models such as BRR and LASSO are the worst performing algorithms, achieving best performance on only one or two datasets.

**Figure 2:**
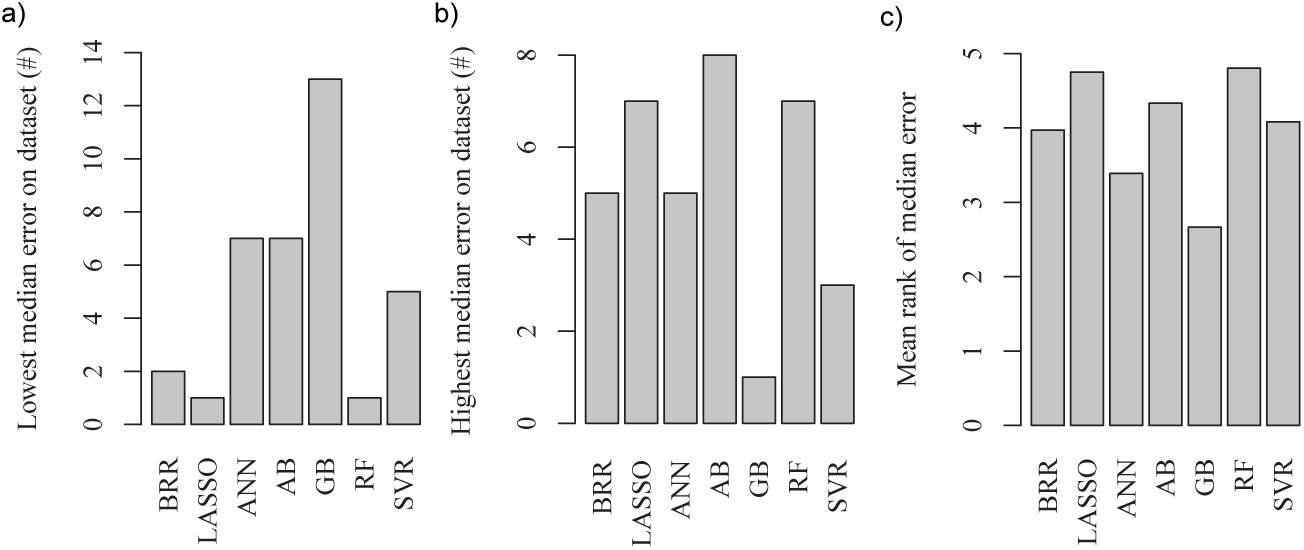
The frequency with which an algorithm scored the lowest (a) and highest (b) absolute median error on a dataset. Panel (c) shows the mean rank over all datasets when the performance is sorted on absolute median error.

Figure 2(b) shows the number of times an algorithm had the highest median absolute error on a dataset. GB yet again performs significantly better than the other algorithms, displaying worst performance for just one dataset (UniToyama Atlantis). The SVR (linear/RBF) model follows closely, having the worst median error on only three datasets. AB is ranked worst for eight out of 36 datasets, while achieving best performance for 5 datasets. This indicates that AB can achieve competitive performance, but is susceptible to overfitting. Overall, this comparison shows that GB will not always perform the best, but out of all algorithms it is most likely to show the best generalization performance, while being least likely to show the worst performance.

Even though there is a clear difference between algorithms regarding best and worst performance, the mean rank for the different algorithms is more similar (Figure 2(c)). This similarity in mean rank indicates that high performing algorithms (GB, MLP, SVR) can at times perform poorly (but not the worst) on some of the datasets. These results show that different algorithms will show different performance on different datasets, thus illustrating the importance of evaluating multiple algorithms for individual datasets.

### 3.3 Pairwise performance comparison of the algorithms

To obtain more insight into the robust performance of GB across the datasets, we compared its performance against the other learning algorithms for each of the 36 datasets. Figure 3 shows the difference in median absolute error normalized to the maximum elution times for each dataset (eqn. 4). GB improves the normalized median error by 0.74 % to 1.51 % on average for all datasets compared to the other six algorithms. For example, in the case of Taguchi a 1 % improvement would translate to a 40.89 second lower median absolute error. The datasets Matsuura, Matsuura 15 and Beck are consistently a worst fit for GB when compared to the other algorithms. For completeness, the pair-wise comparisons are also made for AB, BRR, ANN, Random Forest and SVR models (Figure S-4). The differences in normalized median absolute error range from 0.01 % to 0.77 % and are overall much lower than the differences observed in Figure 3 for GB.

**Figure 3:**
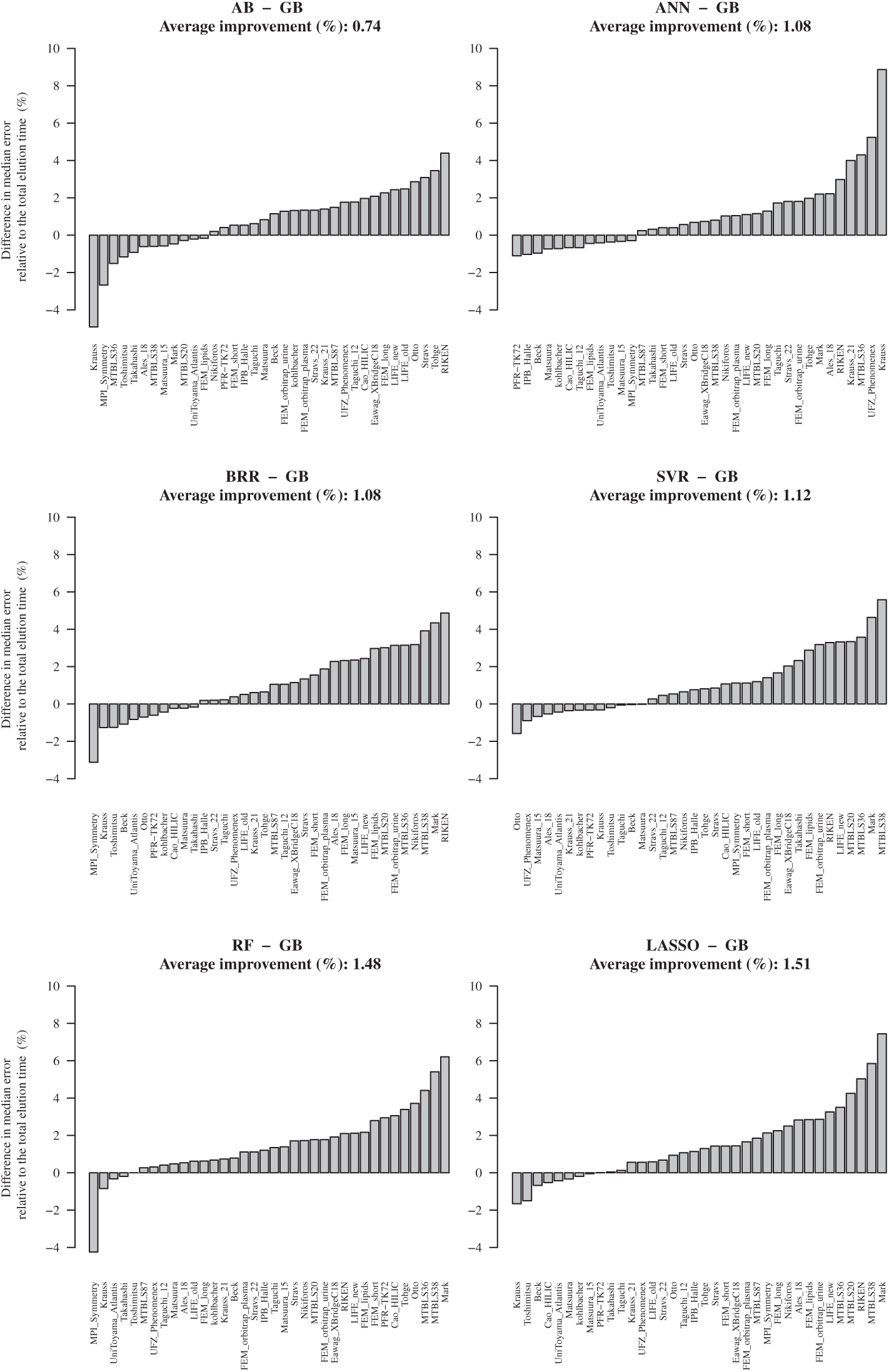
Difference in the absolute median error per dataset for the GB models compared to LASSO, AB, BRR, ANN and SVR. Positive numbers indicate that the median absolute error was lowered (improved) by the indicated amount when using GB. Negative numbers indicate a lower median absolute error for the algorithm GB is compared to.

Figure 4 shows the comparison of choosing the best performing algorithm compared to GB for each dataset. For the thirteen datasets where the GB model was best, the performance increase is, of course, zero. For the remaining datasets, choosing the highest performing model improved the relative median error with 0.72 % on average (Figure 4(a)). Because of its popularity, the same comparison was made for SVR, where choosing the best performing algorithm rather than SVR resulted in an average performance improvement of 1.84 % (Figure 4(b)). This again shows that evaluating multiple algorithms to select the best model can significantly lower the prediction error, and that the popularity of SVR seems unwarranted in light of the overall superior performance of GB.

**Figure 4:**
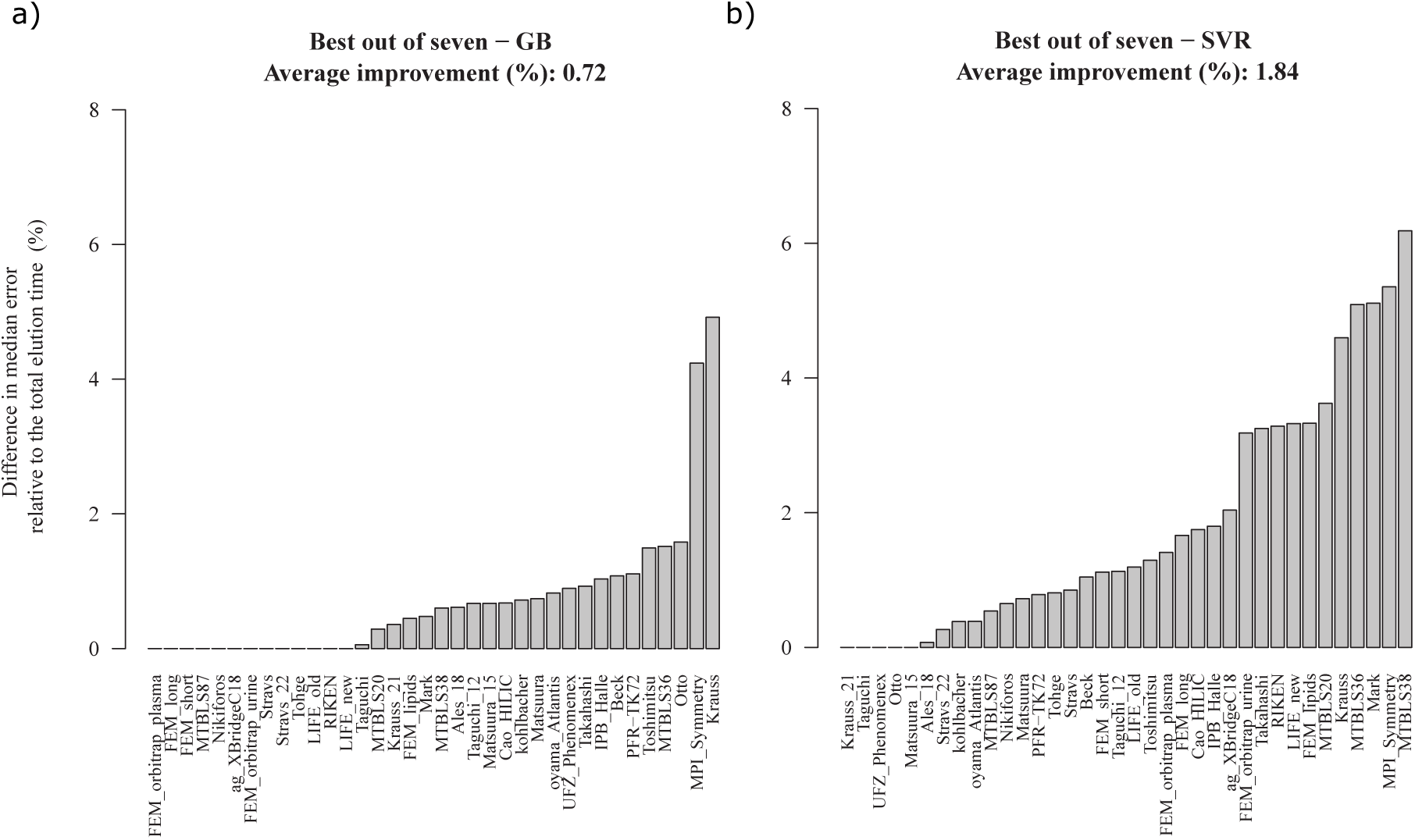
Comparison of the normalized median error between the best performing model out of the selected seven per dataset, and the GB (a), and SVR (b) models. Zero difference in the error indicates that the algorithm under evaluation performed the best. Non-zero values indicate the lower error when choosing one of the other algorithms.

### 3.4 Combining multiple prediction algorithms

Instead of selecting the best learning algorithm for a dataset one might also consider combining the predictions of several models to compute what is called blended predictions. This approach can only achieve better performance than the best performing model alone if the predictions computed by the different models are not too correlated. Figure S-6 shows the correlation between the errors for the different learning algorithms. It shows that this correlation is high (*r* ¿ 0.82) between the three algorithms that fit a continuous mathematical function (SVR, BRR and LASSO), and that it is also high (*r* ¿ 0.73) between the three tree-based algorithms (GB, AB and RF). However, the correlation is much lower between these two classes of three algorithms (*r* ¡ 0.59). Finally, the ANN model shows very low correlation with any of the other algorithms (*r* ¡ 0.58).

A very simple blending strategy was implemented based on these correlations, which averages the predictions of SVR, ANN and GB. While the effect of blending will be minor for molecules that show similar *t*_*R*_ predictions for the three blended models, the potential effect on molecules that show sufficiently different *t*_*R*_ predictions should be the reduction of outlying predictions (i.e. predictions with large error).

This was investigated by looking at the percentage of molecules (over all 36 datasets) with prediction error lower than a certain threshold. Figure 5 shows different values for the threshold on the x-axis with the corresponding percentage of molecules with prediction error below this threshold on the y-axis.

**Figure 5:**
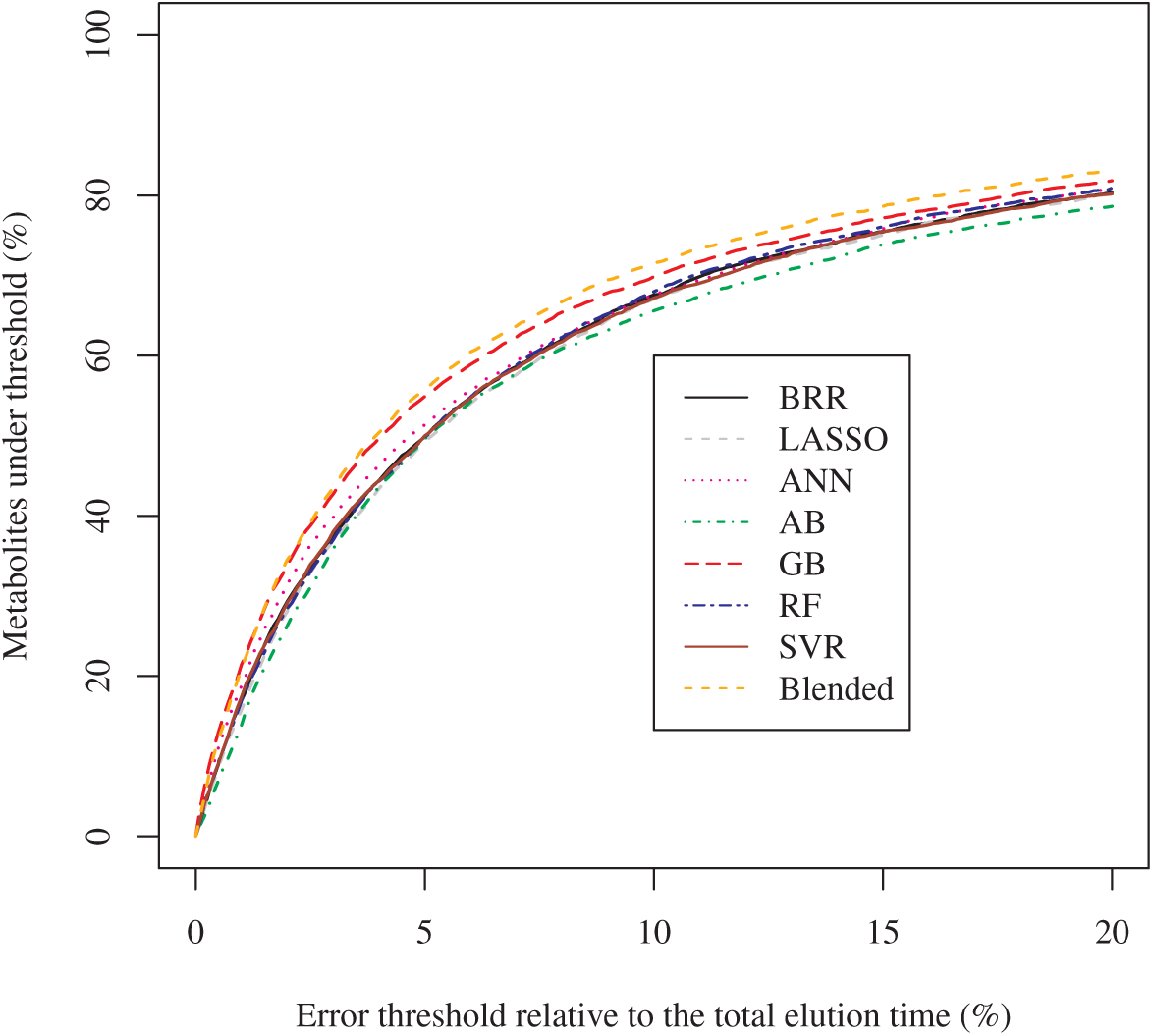
Percentage of metabolites with predictions under an error threshold plotted against that error threshold across all 36 datasets. Blended predictions are calculated as the average prediction for a metabolite by GB, ANN and SVR.

Figure 5 shows that the averaged (blended) predictions perform better than the single best model, except for molecules with the smallest prediction errors (errors ¡ 2.5%). For individual datasets, both the mean and median absolute error rank is improved when blending, and is better than the GB models (Table S-6). So, even though the difference between the blended model and GB is small, these results show that outlying *t*_*R*_ predictions can be reduced by even a simple blending strategy. It is likely that more advanced blending strategies can further reduce prediction error.

However, Figure 5 also shows that only about half of all metabolites are predicted with high accuracy (error ¡ 5%). For instance, for GB the threshold where 50 % of all molecules falls within this threshold occurs at an error threshold of 4.1 %. After this point, higher thresholds provide relatively smaller gains. For instance, for GB the threshold where 80 % of the metabolites falls within this threshold only occurs at an error threshold of 17.7 %.

### 3.5 Comparison between SVR and gradient boosting models

The SVR algorithm is one of the most popular algorithms for *t*_*R*_ prediction, but the results presented here show that the GB algorithm is more accurate for most datasets. The boosting algorithm in GB works by decreasing the bias of an ensemble of weak learners that have a low variance and high bias. The downside of this is that there need to be enough training examples to effectively decrease the bias.

Figure Figure 6 shows a comparison between the performance of both algorithms while taking the size of the dataset into account. The 36 datasets are split into two equal sized groups: one with low, and one with high number of training examples. This division was made based on a threshold of at least 100 training examples for the high number of examples. 13 out of the 18 datasets in the group with a high number of examples have a lower median absolute error for GB than for SVR (Fisher’s exact test p-value *<* 0.02), and 9 out of 18 datasets in the group with a low number of examples have a lower median absolute error for GB than for SVR (Fisher’s exact test p-value = 1).

**Figure 6:**
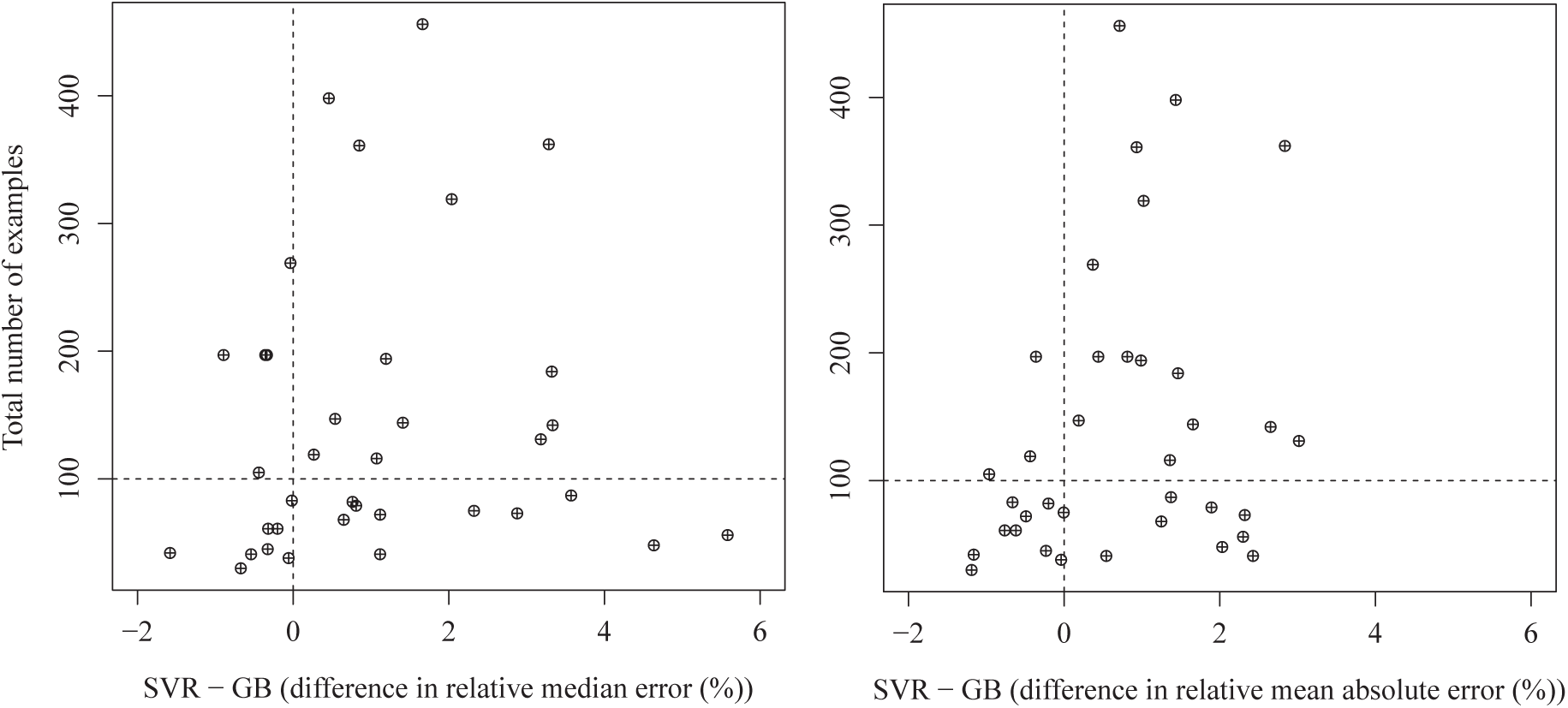
Comparison of the median and mean absolute error between GB and SVR for every dataset. The difference in the median and mean absolute error between GB and SVR is plotted with the number of training examples. The vertical line indicates the area where there is no performance difference, and the horizontal line indicates the 100 examples cut-off between low and high number of examples in the dataset.

The mean absolute error shows the same trend, where 15 out of 18 datasets with high number of examples have a better performance for the GB trained model (Fisher’s exact test p-value *<* 0.0002), while 10 out of 18 datasets with low number of examples have a better performance for GB (Fisher’s exact test p-value = 0.74).

### 3.6 Effect of a reduced feature set

In this section we want to investigate the possible overfitting by the machine learning algorithms. Overfitting is a problem for any machine learning task, but due to the relatively high amount of features (151) compared to the number of training examples this becomes a serious concern for some datasets. To detect overfitting on 151 features, the performance of models trained on all 151 features is compared with models trained on the minimal subset of eleven features. This because overfitting is less likely to occur in the latter.

Figure 7 shows that the three datasets (Matsuura, Matsuura 15 and Beck) that were consistently a worse fit for GB in Figure 3 achieve a higher performance on the minimal set of eleven features. This higher performance for the minimal feature set indicates overfitting on the 151 features. However, limiting the number of features decreased the performance for most datasets (23 out of 36) with on average a lower median absolute error of 0.42 % for GB models trained on all 151 features. This shows that the larger feature set is still preferred over the minimal feature set for most datasets.

**Figure 7:**
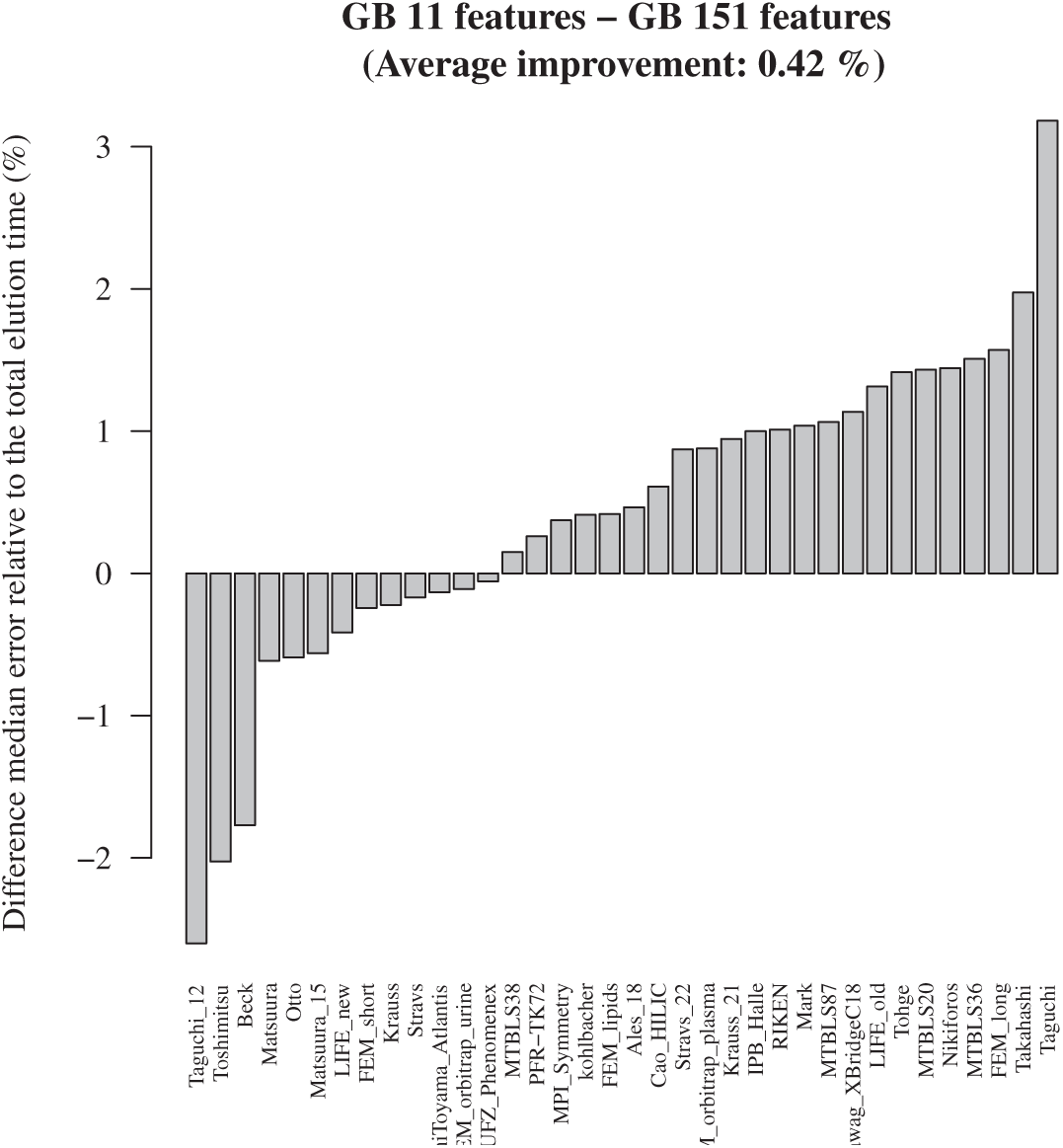
Difference in the absolute median error per dataset for GB trained with eleven, or with 151 features. Positive numbers mean that the absolute median error was lowered by the indicated amount when applying GB with 151 features.

The pairwise comparison between the other learning algorithms that are fitted on the reduced feature set in Figure S-5 follows the same conclusions as Figure 3. GB is able to outperform the other algorithms with an improved median absolute error of 0.94 % to 2.75 % on average. This shows that the earlier conclusions made with the complete feature set are not due to overfitting of some of the algorithms on the large set of features.

### 3.7 Feature relevance

The performance evaluations conducted here have shown that overall, GB models perform best. The relevance of each feature in all GB models is therefore investigated in more detail in this section. The F-score provided by GB reflects this feature relevance, and is computed as the number of times that feature was selected to split the training examples during decision tree construction.

Figure 8 shows the highest scoring feature based on the F-score. The MolLogP describes the hydrophobicity and is the most important feature across all datasets. This is the result of the large proportion of datasets based on reverse-phase LC, which separates analytes based on hydrophobicity. The remaining features generally describe structural features of molecules on top of chemical features (Chi4v, kappa, EState, PEOE and SlogP). The relevance of these features can be explained by interactions of reactive groups (e.g. cyclic carbons) with the solid phase of the column, and similarity between molecules. High structural and chemical similarity between molecules is therefore a good indicator for molecules to have the same retention time.

**Figure 8:**
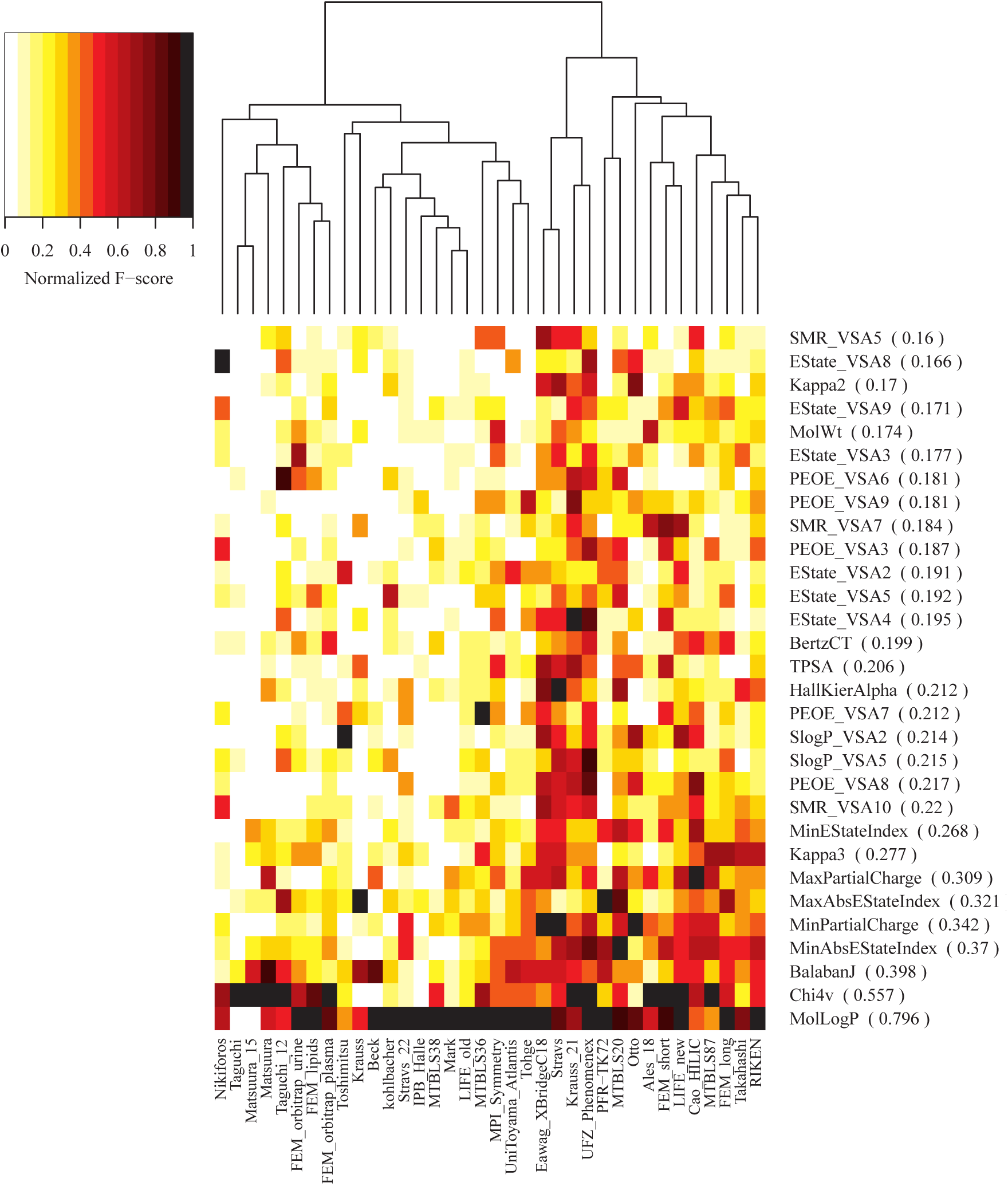
Normalized F-score from GB per dataset. F-scores were normalized by dividing all features by the maximum F-score per dataset. Every feature name is followed by the mean F-score of all datasets. Any feature with a mean F-score below 0.15 is excluded.

Figure 8 also shows that the importance of features is often shared between datasets, but that a substantial proportion of features used in the models is nevertheless shared by only a few datasets. These differences in feature importance underline that knowledge of features cannot be directly transferred from one experimental setup to another.

## 4 Conclusion

When making an LC *t*_*R*_ predictor, researchers make multiple decisions that can influence the performance and interpretability of the final model. The most notable such decisions are the number of training examples to include, the machine learning algorithm to use, and the molecular features to calculate.

For all datasets analysed here, a fairly accurate model could only be obtained when the number of training examples was at least 40. However, different datasets and algorithms require a different number of training examples to achieve the highest performance possible. Generally the GB algorithm delivers the best performance, given sufficient training examples (¿100). GB models generally keep improving for increasing number of training instances, where other algorithms converge in their performance more quickly as training instances increase.

Cross-validation confirms that GB is the most likely candidate to deliver the best performance, and is the least likely to give the worst performance. However, the best performance in 23 out of 36 datasets was obtained using another algorithm. This shows the importance of testing different algorithms before choosing the learning algorithm that is used to generate the final model. GB, ANN and SVR generate complementary models and provide a good starting point for blending multiple algorithms.

The feature importance analysis of the GB models show that feature relevance is also dataset dependent. Selecting features based on other experimental setups can therefore result in a suboptimal model. In the initial stages of creating a model, a researcher should therefore be careful when excluding features. GB is generally able to select those features that are important to achieve a high performance without overfitting.

However, the best overall performance can be achieved by blending algorithms, which results in a lower overall prediction error. The blending technique applied here is very simplistic, but this already indicates that more advanced blending techniques are worth investigating.

## 5 acknowledgement

This project was made possible by MASSTRPLAN. MASSTRPLAN received funding from the Marie Sklodowska-Curie EU Framework for Research and Innovation Horizon 2020, under Grant Agreement No. 675132.

